# Impact of mixing between parallel year groups on genomic prediction in Atlantic salmon breeding programmes under random selection

**DOI:** 10.1101/2022.12.12.520109

**Authors:** Panagiotis Kokkinias, Alastair Hamilton, Ross Houston, Chris Haley, Ricardo Pong-Wong, Pau Navarro

## Abstract

A commercial breeding programme in Atlantic salmon utilises a four-year generation interval with four parallel breeding populations. In this study, we develop a computer simulation of a salmon breeding programme and explore the impact of gene flow between the parallel year groups on the accuracy of genomic prediction within and between breeding lines. We simulated four parallel lines for 10 discrete generations with random selection and different mixing rates between parallel year groups. The genetic distance between fish (as a measure of diversity) and the accuracy of estimated genomic breeding values were used as criteria of comparison. With no mixing the genetic distance increased between populations, the genetic variation within populations decreased and there was no increase in accuracy when combining data across populations. Even a low percentage of mixing decreased the genetic distance between populations and increased the genetic variation within populations. The higher the percentage of mixing the faster the lines became more similar. The accuracy of prediction climbed as the percentage of mixing increased. The increase in accuracy from the combined evaluation approach compared to the within evaluation approach was greater with an increased percentage of mixing. In conclusion, if there is no gene flow between populations the lines drift apart and there is no value in combining information across populations for genomic breeding value prediction. Only a low amount of mixing between lines brings the lines closer together and facilitates the use of information across lines to improve breeding value prediction. Optimising gene flow between lines should be an integral part of salmon breeding programme design.

## 1. Introduction

Improved stocks of aquaculture species with better utilization of feed, land and water resources are critical for the development of a profitable commercial production (Troell et al. 2014). Genomic prediction methods are rapidly becoming important selection tools in livestock breeding as they have the potential to substantially increase the accuracy of the estimated breeding values (Meuwissen, Hayes and Goddard 2001). Recent studies in aquaculture species have shown that genomic selection can achieve higher prediction accuracy of breeding values than the traditional pedigree information alone, especially within families (Meuwissen et al. 2014, Ødegård et al. 2014, Tsai et al. 2015). However, despite the theoretical benefits of these methods, in practice increases in accuracy are lower than those predicted by the theory when considering distant relatives or across different populations (Kachman et al. 2013).

Atlantic salmon experience a major part of their life and growth in salt water and migrate back to freshwater as adults to spawn and complete their life cycle. In farmed conditions, during the first-year fertilized eggs hatch and juveniles grow through several distinct stages in freshwater. As a result of the four-year lifecycle of salmon in Scotland and Norway, aquaculture-breeding programmes tend to be composed of four parallel and isolated sub-populations (“Year Cycles” or “Lines”) with one-year difference between them to provide stock for the farming industry each year. Normally, broodstock parents are not reused across generations and therefore the generations are discrete (i.e. they do not overlap) (Gjedrem et al. 1991).

Although the four lines in a breeding programme generally have the same origin (Gjedrem et al. 1991), their isolation over a number of years of production leads to genetic differentiation between them. This differentiation leads to increasing genetic distance between the populations that can compromise the efficacy of predicting genomic estimated breeding values (GEBV) for individuals of a given line using data from other lines. This genetic differentiation creates the need for separate genetic evaluation within each line. The consequence is genomic evaluation with a smaller training population reducing the accuracy of the GEBV, thereby reducing genetic gain (Gjøen and Bentsen 1997).

Recently, the aquaculture industry has introduced the production of “underyearling” smolts by accelerating their growth during their first months. This allows transfer of smolts to sea water prior to the onset of winter and accelerates sexual maturation of the broodstock, reducing the production cycle to three years (Duston and Saunders 1995). In order to increase the connectedness between the lines, sexually mature three-year-old individuals from a line could be used as parents a year earlier (i.e., at three rather than four years) in another line (this will be referred as “mixing”). This movement of individuals across the breeding lines should decrease the genetic distance between them, so data from other lines may provide useful information in a joint genetic evaluation creating a larger reference population and improving the accuracy of the genomic evaluation.

The objective of this study was to investigate using computer simulation the impact of the rate of mixing on the accuracy of prediction from genomic evaluation in a typical Atlantic salmon breeding programme. The simulations were based on four parallel populations with simulated phenotypic and genotypic data over 10 breeding cycles.

## 2. Materials and Methods

### 2.1. Simulation of the base population in linkage disequilibrium for each line

The base population of each line in the breeding programme was simulated in three steps: (1) creation of a founder population in linkage disequilibrium; (2) differentiation of the lines; and (3) expansion of the population. The full protocol for generating the base population for each line is shown in Figure 1.

**Figure 1.**
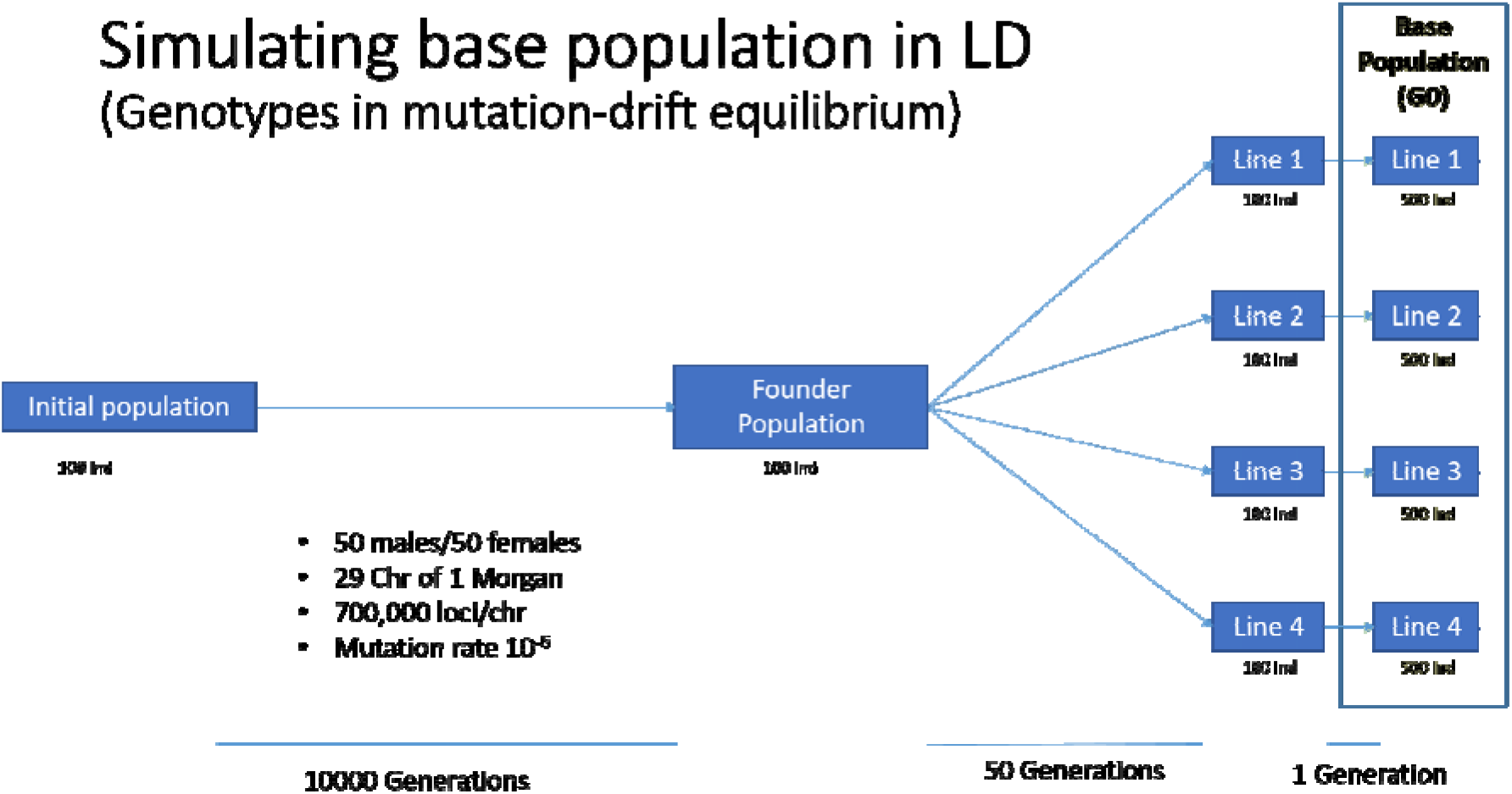
Steps of simulation from initial population to base population (G0). Ind: individuals, Chr: chromosome.

A founder population in linkage disequilibrium (LD) was simulated using a Fisher-Wright population model (Fisher 1930, Wright 1931, Sonesson and Meuwissen 2009). Briefly, an initial population of *N* individuals is allowed to reproduce by mating at random, with each individual producing two offspring (one male and one female). Their genome is composed of several chromosomes with biallelic loci mutating at a given rate. As the population develops across a large number of generations, new mutations appear. These mutations increased or decreased in frequency due to genetic drift, resulting in a population with LD between closely linked loci. For this study, the initial population was composed of 100 individuals (50 males and 50 females). The genome consisted of 29 chromosomes of 1 Morgan, each with 700,000 loci with mutation rate set at 10^-6^. After 10,000 generations, around 3,100 loci were segregating in each chromosome (approximately 90K across the whole genome). Individuals at generation 10,000 are denoted as the founder population.

In the second step the breeding lines were simulated. Here, the individuals of the founder population (i.e., individuals from generation 10,000) were randomly mated for a further 50 generations, using the same algorithm as before but no mutation was allowed. Each line was created by repeating the same protocol independently starting from the same founder population. This resulted in the creation of four lines with the same origin but with some degree of divergence due to drift. Thereafter, at the third step, individuals from generation 10 (50 males and 50 females within each line) were mated randomly to create 500 progeny which were considered as the base population (G0) for each line of the breeding programme. The simulation analyses were performed using software developed in-house.

### 2.2. Structure of the salmon breeding population

A commercial Atlantic salmon breeding programme, with a generation interval of four years (one breeding cycle), was simulated for 10 cycles. In order to have a harvest every year, each nucleus breeding programme consisted of four sub-population (breeding lines) which are isolated, with discrete cycles and candidates (50 males and 50 females) that cannot be reused (blue arrows in Figure 2).

**Figure 2.**
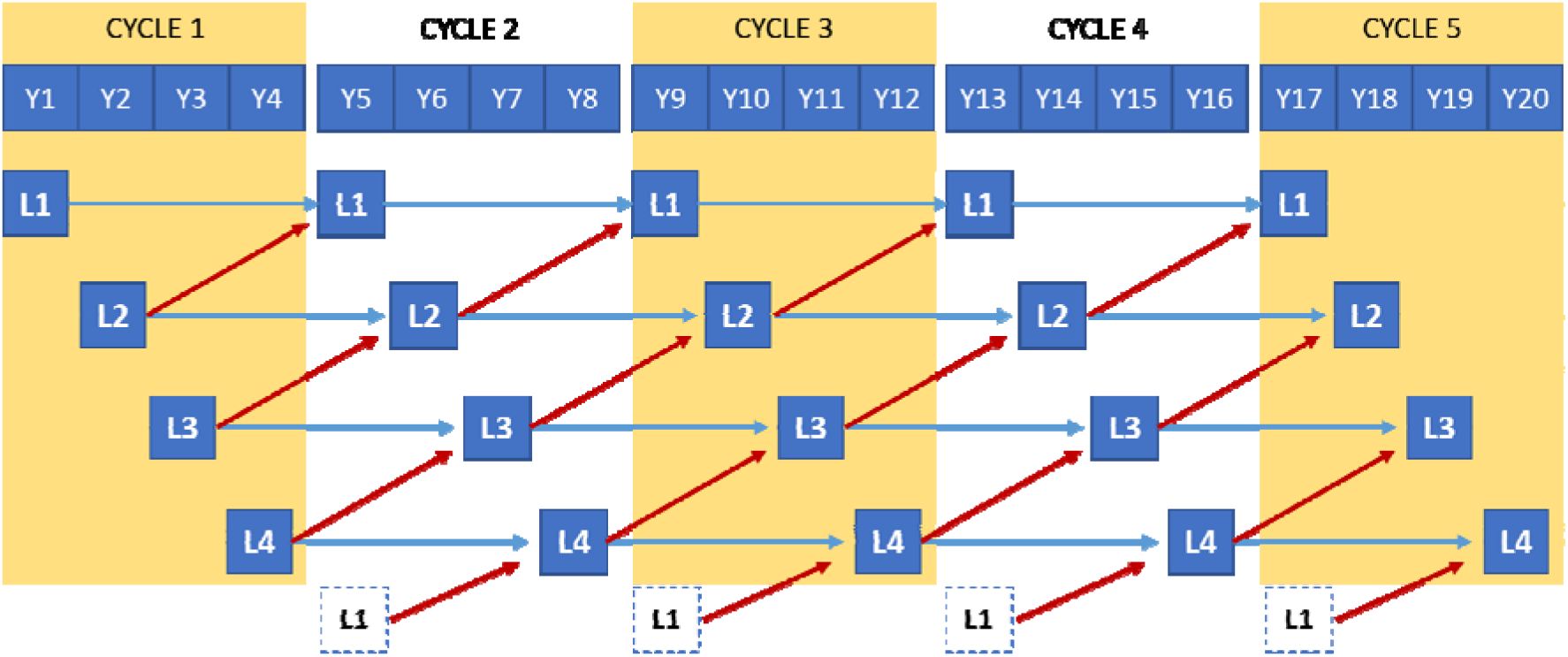
Four parallel breeding populations with a four-year cycle interval. Blue arrows show the gene flow within a line and the red arrows between the lines by using underyearling individuals (mixing). Example of the first five cycles.

To increase the connectedness between the lines, we introduce a proportion of individuals from one line to another (“mix”). Some individuals mature at three years and can be used as parents. For example, individuals from line 1 and underyearling individuals from line 2 will be used to create the next cycle of line 1. Different mixing rates (percentage X% of 3-year-old candidates) have been tested to investigate the effect of the mixing rate on the accuracy of GEBV (Figure 2). For example, with 10% mixing rate 90 individuals (45 males and 45 females) from line 1 and 10 individuals (5 males and 5 females) from line 2 will be used as candidates to create the next breeding cycle of line 1.

### 2.3. Genetic architecture

In each chromosome 100 segregating SNPs were randomly sampled to be used as QTL controlling the trait, and 1000 SNPs were randomly selected to be used as genetic markers in the genomic evaluation (i.e., in total across the 29 chromosomes, 2900 loci were selected as QTL and 29000 as SNPs in chip (so were “genotyped”) with no overlap between the QTL and chip SNPs). The trait was simulated to have a heritability of 0.2 and a phenotypic variance of 100.

The simulated true breeding value (TBV) for an individual was calculated as:where *m* is the total number of QTL, *g*_ij_ is the genotype score of the QTL *j* for individual *i*, coded as 0,1,2 when the genotype is AA, AB and BB, respectively.

The genotype score is the number of the reference alleles (B) in the genotype and *a_j_* is the additive effect for QTL_*j*_. The additive effects were sampled from a normal distribution and rescaled such as the variance of the TBVs in the base population (i.e., G0) was 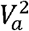.

The phenotypic value, *y*_i_ of individual *i* was obtained by adding a normally distributed environment effect E_i_, (with mean to zero and variance 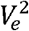) and a year effect Y_eff_ to the genetic value (TBV) equal to:

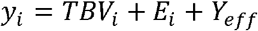

The year effect was sampled as a random variable, uniformly distributed between 0 and 10, which is equivalent to a range of 1 phenotypic standard deviation.

### 2.4. Genomic estimated breeding values

The genomic estimated breeding values (GEBV) were calculated based on the genomic best linear unbiased prediction (GBLUP) method by VanRaden (VanRaden 2008) for all the individuals. The following mixed model was used to estimate the GEBV:

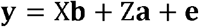

where **y** is the vector of observations, X the design matrix for the vector of fixed effects **b** (e.g., year cycle effect), Z is the incidence matrix that relates animal effects to the individual records, **a** is the vector of animal effects with size **n** (the number of individuals) and **e** is the vector of random residuals following a normal distribution N (0, I *V*_e_^2^), in which *V*_e_^2^ is the variance of residuals. The animal effects follow a normal distribution N (0, G *V*_e_^2^). The matrix G is the genomic relationship matrix calculated using the VanRaden algorithm as follows:

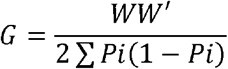

Where W is a centralised genotype matrix with rows as individuals and columns as markers. *P_i_* is the frequency of the second allele for the marker *i*. For each analysis a new genomic relationship matrix was estimated, and the *P_i_* values were recalculated, rather than re-using elements from a big across-population matrix. This genomic relationship matrix is derived from allele frequencies as suggested by VanRaden and the evaluation was done with animals from the same cycle (VanRaden 2008).

In order to explore different scenarios, the above model was applied by using data in the three following ways:

1. Using data from a population (line) to calculate the GEBVs for the individuals of that population (same line), referred as GBLUP-W.
2. Using data from one population (e.g., line 1) to calculate the GEBVs for the individuals of another population (e.g., line 2) with unknown data (phenotypes), referred as GBLUP-B.
3. Using all data (4 lines) jointly to calculate GEBVs of individuals of one population (e.g., line 1), referred as GBLUP-C.

### 2.5. Scenarios compared

Seven scenarios were compared: one where there is no mixing between the lines and 6 scenarios where there is a proportion of mixing between the lines. The mixing rate for these scenarios were: 4, 10, 20, 30, 40 and 50%. These different mixing rates were used in order to investigate how the genetic distance and the accuracy of GEBVs change during the 10 cycles as we increase the gene flow between the breeding lines.

### 2.6. Criteria of comparison

In order to explore the impact of different mixing rates between populations, we use two criteria of comparison between the different scenarios: the genetic distance between the individuals and the accuracy of estimated genomic breeding values. The results were based on 10 replicates for each tested rate of mixing and the average of the replicates was reported.

#### Genetic distance

To calculate the genetic distance within and between the populations in each cycle, a genomic relationship matrix (GRM), that includes all the individuals, was calculated at each cycle. An Eigen decomposition was done on the GRM to obtain its Eigenvalues and Eigenvectors. Since the Eigenvectors are orthogonal among themselves, they can be used to calculate the Euclidean distance between two individuals.

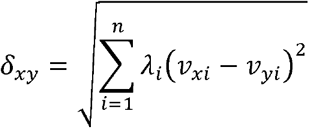

where *v_xi_, v_yi_* is the value of the eigenvector *i* for animals x and y respectively, λ_*i*_ the eigenvalue *i* and **n** is the total number of the eigenvectors with non-zero associated eigenvalue.

The genetic distance is the average of all pairwise Euclidean distances between individuals of the same or different line. The average between individuals of same line is referred as “Within” and used only data (pairwise Euclidean distances) of that line. The average of individuals that belong to consecutive lines (i.e. with only one year difference and/or direct gene flow because of mixing) is referred to as “Consecutive” and used only data from the lines in question, while the average between individuals that belong to non-consecutive lines (i.e. with two years difference and no direct gene flow between them) is referred to as “Non-Consecutive” (Figure 3).

**Figure 3.**
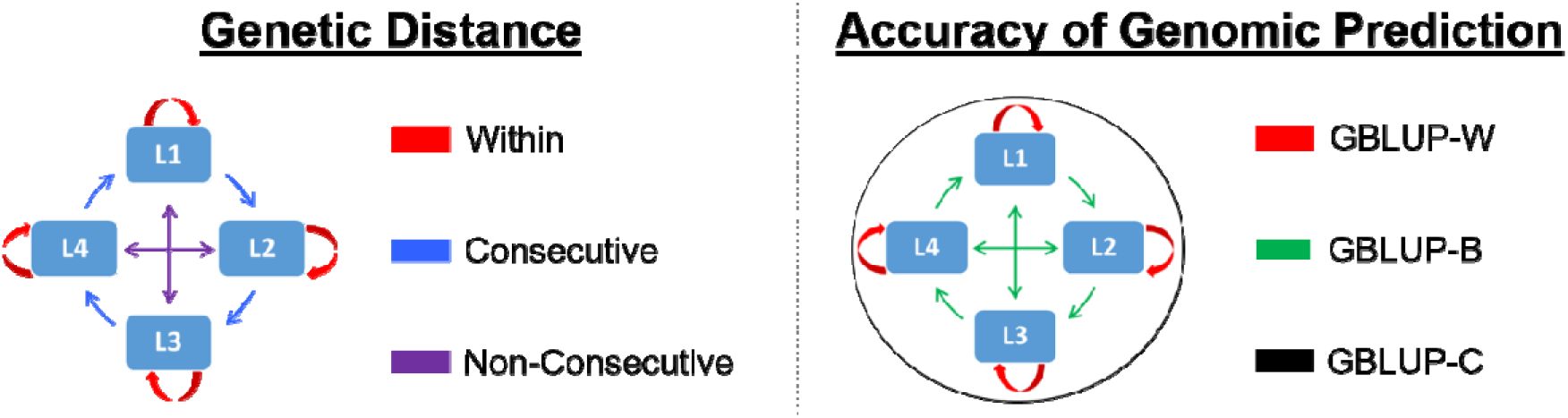
Description of the different genetic distances and accuracies of genomic prediction estimates compared in our study. The average genetic distance between individuals of same line is referred as “Within”. The average of individuals that belong to consecutive lines is referred to as “Consecutive” while the average between individuals that belong to non-consecutive lines is referred to as “Non-Consecutive”. The accuracy of the genomic was estimated based on three different schemes: A) Using data from only one line to estimate the GEBVs of the same line (GBLUP-W), B) using data of two lines where in one line the data are known and we estimate the accuracy of predicting GEBV of the other line (GBLUP-B) and C) using data from all the populations jointly to estimate the accuracy of predicting GEBVs of one line (GBLUP-C).

#### Accuracy of genomic prediction

The accuracy of the genomic prediction is defined as the correlation between the true breeding values (TBV) and those estimated (GEBV). In each replicate, the accuracy was estimated based on three different schemes: A) Using data from only one line to estimate the GEBVs of the same line. The average accuracy of the four lines referred as GBLUP-W, B) using data of two lines where in one line the data (TBV and GEBV, e.g., line 1) are known and we estimate the accuracy of predicting GEBV of the other line (e.g., line 2, where the GEBVs are unknown). The average accuracy (of all the possible combinations between the lines) is referred as GBLUP-B and C) using data from all the populations jointly (merging all lines and using as one population) to estimate the accuracy of predicting GEBVs of one line (e.g., line 1). The average of the four values (one estimate of accuracy for each line) is referred as GBLUP-C (Figure 3).

The evaluation of the combined data (GBLUP-C) will allow to estimate the impact of using a combined reference population (hence increasing the training population as it uses jointly the four lines) and compare it with estimates when using data from the same line (GBLUP-W) and from a different line (GBLUP-B) as training data.

## 3. Results

### 3.1. No mixing scenario

Four discrete populations (lines) were simulated and studied for 10 cycles to test how the genetic distance and the accuracy of GEBV change when there is no mixing of individuals between the lines (Table A1, Appendix).

The average genetic distance, between all pairs of individuals within a line (“Within”) and between consecutive (“Consecutive”) and non-consecutive lines (“Non-Consecutive”), was calculated for 10 cycles with no mixing between the lines. The results show that the average genetic distance slightly increased between populations and decreased within populations after 10 cycles. The average distance between populations (Consecutive and non-Consecutive) started with a value of ≈ 1.50 and increased to 1.52 after 10 cycles, whereas the average distance within the lines started at 1.35 and declined to 1.32 at cycle 10 (Figure 4). Through the 10 breeding cycles, the consecutive and non-consecutive line results are very similar and hence overlap with each other, so there is not an obvious separation between them. Moreover, the four lines are very similar through the beginning of the breeding programme (“Within” genetic distance until cycle 6) but there are signs of separation after 10 cycles, which indicate a differentiation probably because of their isolation (Figure 4).

**Figure 4.**
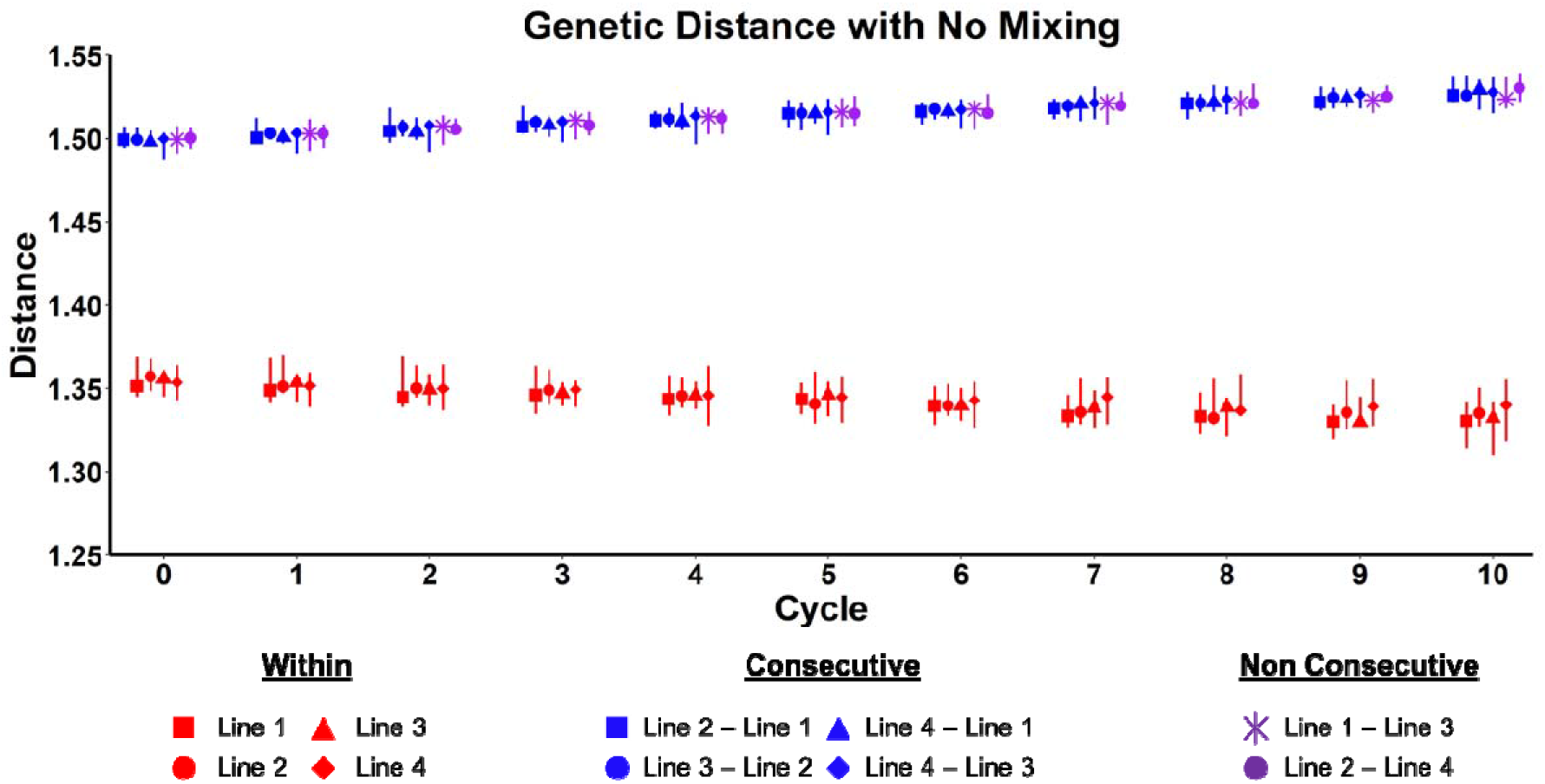
Genetic distance within and between the breeding lines with no mixing for 10 cycles. The points indicate the overall average value and the vertical bars the range of values of the 10 replicates. Average genetic distances are presented for all pairs of individuals within a line (“Within”), between consecutive (“Consecutive”) lines and non-consecutive lines (“Non-Consecutive”).

The average accuracy of genomic prediction, for the 10 replicates, was calculated under three different schemes as described above (Materials and Methods, Figure 3). The GBLUP-W and GBLUP-C accuracy increased through the breeding programme and at cycle 10 they had increased by 8.05% (from 0.641 in C1 to 0.698 in C10) and 8.30% (from 0.644 in C1 to 0.702 in C10) respectively but there was not a significant difference in accuracy between the GBLUP-W and GBLUP-C schemes. The accuracy of the GBLUP-B scheme remained very low, close to zero through the 10 cycles (Figure 5).

**Figure 5.**
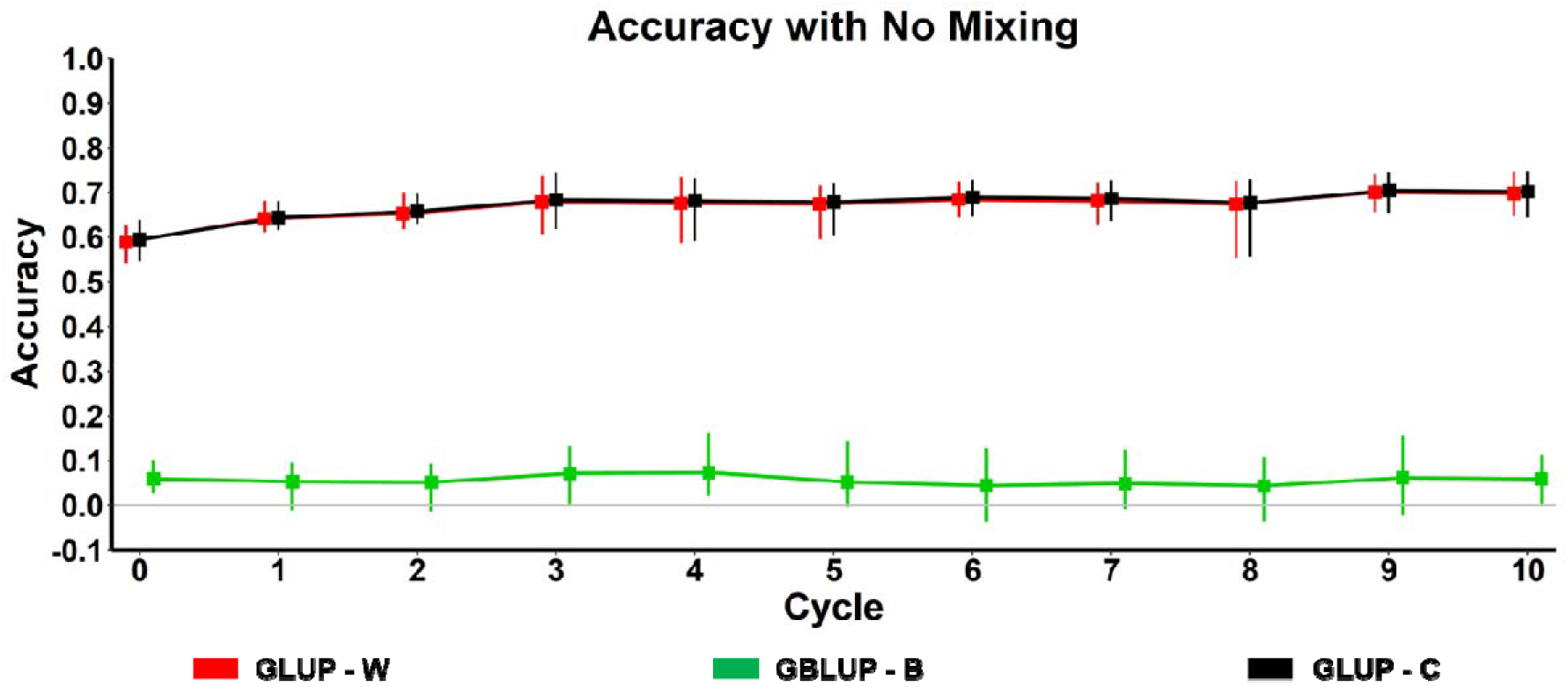
Accuracy of genomic prediction with no mixing for 10 cycles. The points indicate the overall average value and the vertical bars the range of values of the 10 replicates. Results are presented for three different schemes: A) Using data from only one line to estimate the GEBVs of the same line (GBLUP-W), B) using data of two lines where in one line the data are known and we estimate the accuracy of predicting GEBV of the other line (GBLUP-B) and C) using data from all the populations jointly to estimate the accuracy of predicting GEBVs of one line (GBLUP-C).

### 3.2. Scenarios with different mixing rates

Several rates (4, 10, 20, 30, 40 and 50%) of mixing individuals between the four lines were simulated and studied for 10 cycles. Different mixing rates were used to test how different gene flows affect genetic distance and accuracy of prediction.

The average genetic distance was calculated for each mixing rate scenario between individuals of the same line (“Within”) and between individuals of different lines (“Consecutive” and “Non-consecutive”) (Figure 6). A low mixing rate (4%) increased the genetic distance (i.e., the genetic diversity) within each line and decreased the genetic distance between the lines through the 10 cycles. The decrease is smaller between the non-consecutive lines. As the mixing rate increases the distance between the lines (consecutive and non-consecutive) reduces and the distance between individuals within each line (i.e., within line diversity) increases. For 30% mixing, at cycle 5, the average genetic distance within the lines is very similar to the distance between consecutive lines and they overlap at cycle 10. With a mixing rate of 50%, we observe that after only one cycle the distance within lines is very similar to the distance between the lines. At cycle 3, there is no difference between consecutive and non-consecutive lines and the distance within and between lines overlaps. Moreover, the range of values of the 10 replicates (vertical lines) is much wider with a lower than with a higher mixing rate after 10 cycles.

**Figure 6.**
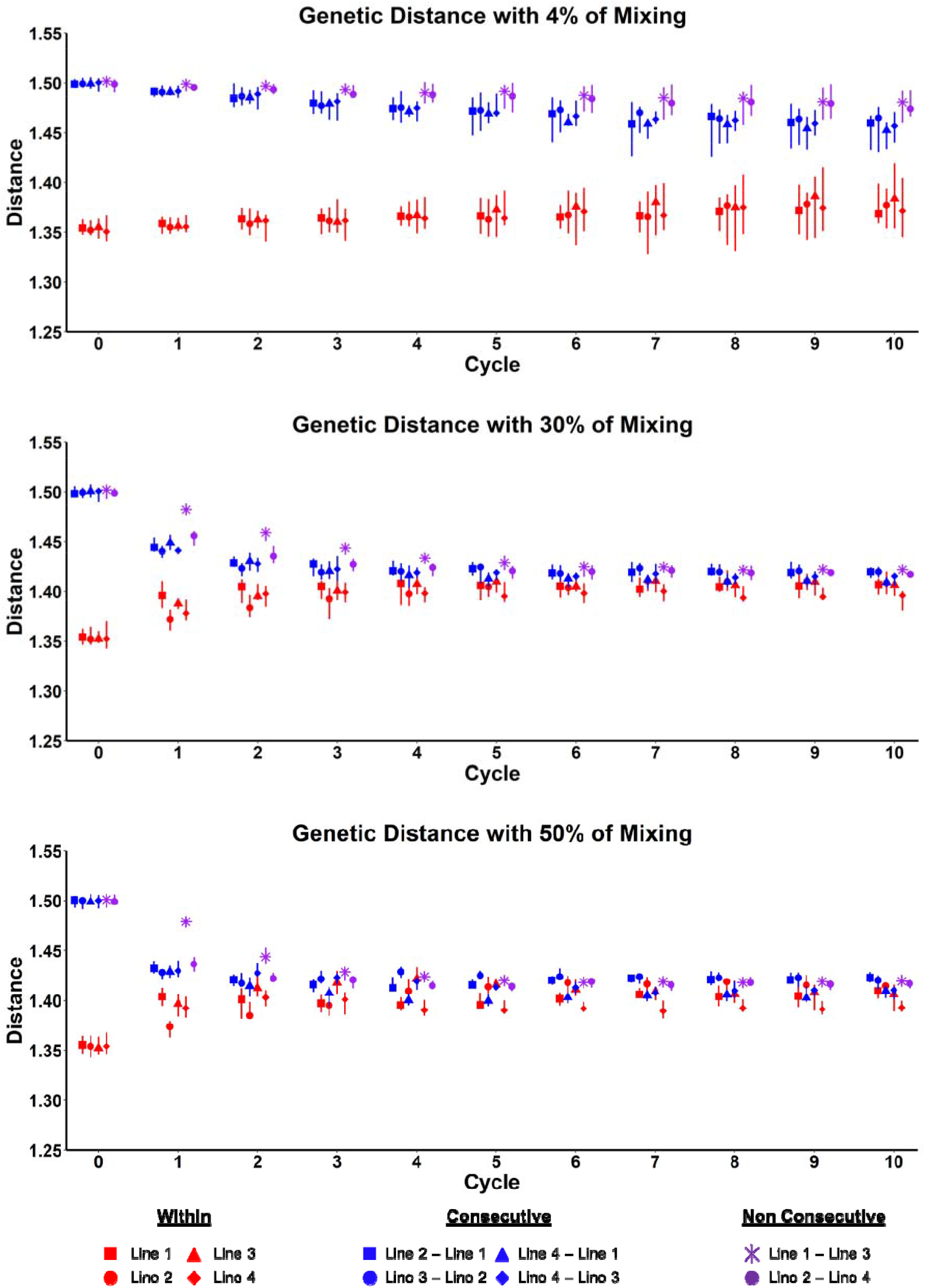
Genetic distance within and between the breeding lines with different mixing rates for 10 cycles. The points indicate the overall average value and the vertical bars the range of values of the 10 replicates. Average genetic distances are presented for all pairs of individuals within a line (“Within”), between consecutive (“Consecutive”) lines and non-consecutive lines (“Non-Consecutive”).

The accuracy of prediction for each scheme (GBLUP-W, GBLUP-B and GBLUP-C) calculated in each cycle under different mixing rates (4, 10, 20, 30, 40 and 50%) is presented in Figure 7 and Table A1 (Appendix). With a low mixing rate (4%), the GBLUP-B accuracy increased by 38.21% (from 0.08 in C1 to 0.13 in C10), the accuracy of GBLUP-W and GBLUP-C increased also by 3.07% (from 0.66 to 0.68) and 5.72% (from 0.67 to 0.72) after 10 cycles.

**Figure 7.**
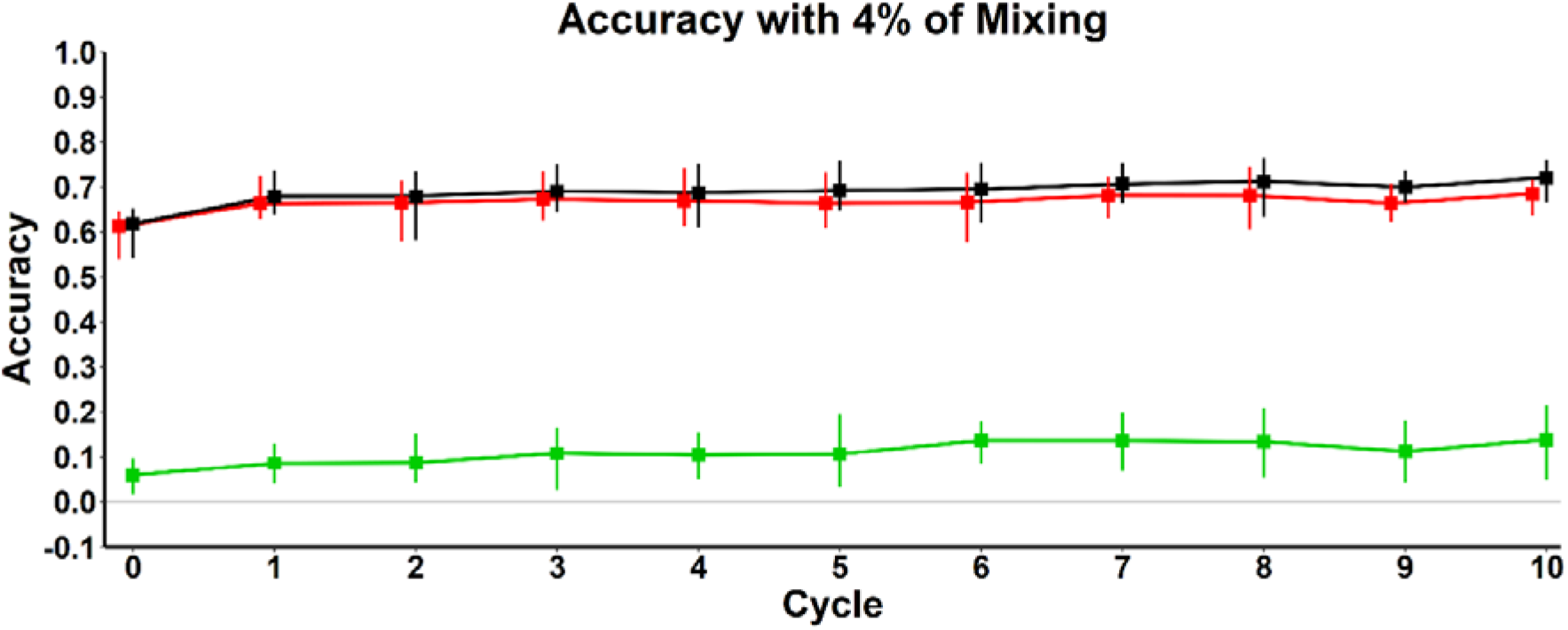
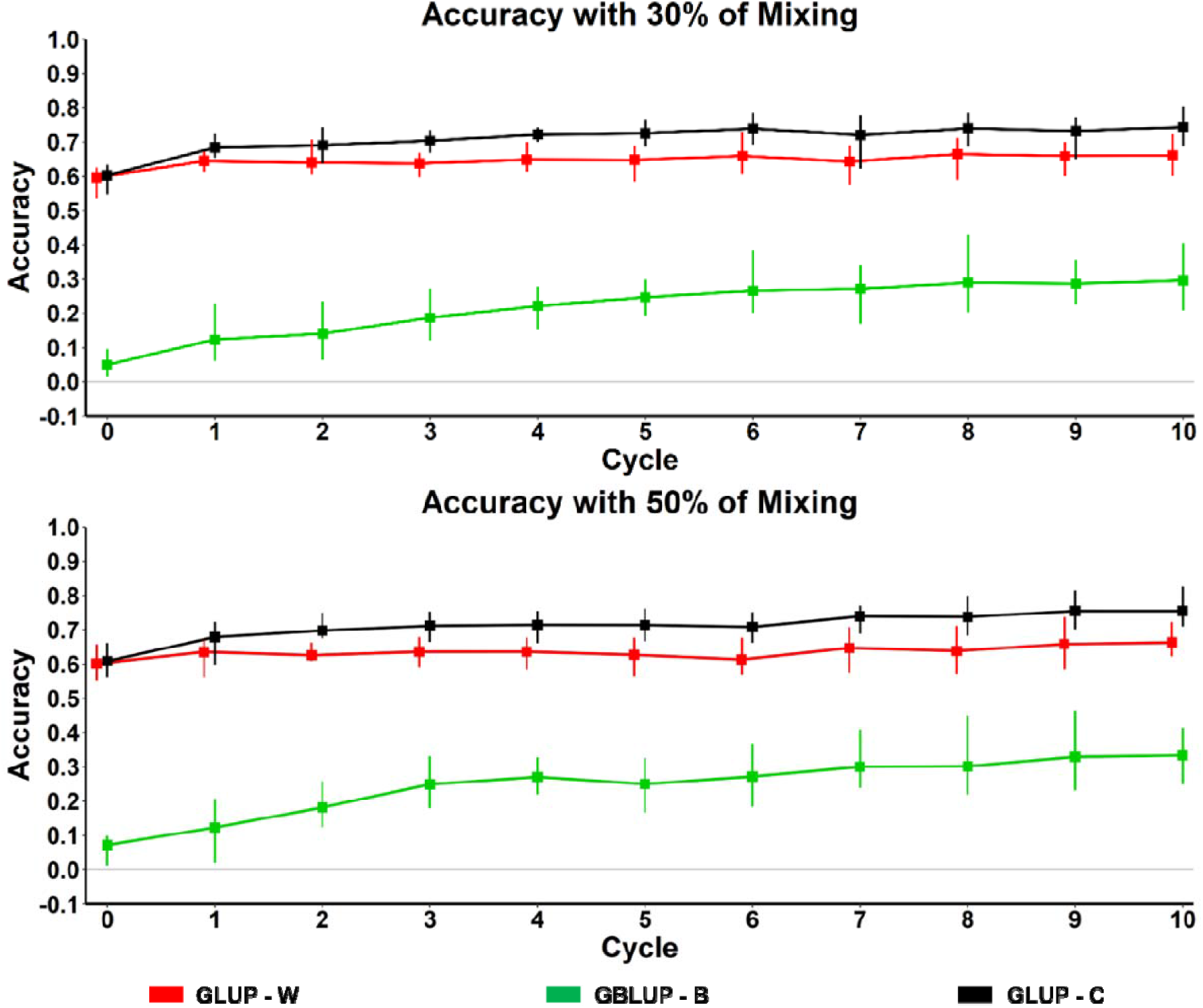
Accuracy of genomic prediction with different mixing rates for 10 cycles. The points indicate the overall average value and the vertical bars the range of values of the 10 replicates. Results are presented for three different schemes: A) Using data from only one line to estimate the GEBVs of the same line (GBLUP-W), B) using data of two lines where in one line the data are known and we estimate the accuracy of predicting GEBV of the other line (GBLUP-B) and C) using data from all the populations jointly to estimate the accuracy of predicting GEBVs of one line (GBLUP-C).

As the percentage of mixing increased, the accuracy of GBLUP-B increased at a higher rate than that of GBLUP-W and GBLUP-C. For example, with a 30% mixing rate, the accuracy of GBLUP-B after 10 cycles had increased by 58.27% (from 0.12 to 0.29), that of GBLUP-W by 2.37% (from 0.64 to 0.66), and that of GBLUP-C by 8% (from 0.68 to 0.74). Moreover, with a mixing rate of 50% the GBLUP-B accuracy increased by 63.08%, that of GBLUP-W by 4.11% and that of GBLUP-C by 10.10% after 10 cycles.

The trend in accuracy of each scheme across the different mixing rates through the 10 cycles is determined by the data used by each model (Figure A1, Appendix). We observe that the accuracy of GBLUP-B increases as the mixing rate increases and achieves the highest increase in rate with mixing between the schemes. GBLUP-W increases for cycle 1 but remains similar through time and achieves higher values with no mixing or low mixing rates. The values of GBLUP-C increase after each cycle for all mixing rates and this scheme provides greater accuracy compared to GBLUP-W (Figure A1, Appendix).

## 4. Discussion

In this study, we investigate by simulation the impact of various mixing rates on the genetic distance and the accuracy of genomic prediction between the four discrete lines of an aquaculture breeding programme. The mating structure and the genetic diversity between the four populations of the base generation (G0) mimics a typical Atlantic salmon breeding scheme. Thus, the main objective was to investigate how much the accuracy of genomic prediction can be increased by using data from combined populations instead of using data from only individual populations to evaluate the same or a different population of the breeding programme.

Our results show that with no mixing, throughout the programme, the genetic distance increased between the lines and decreased within the lines (Figure 4). Hence, the genetic diversity within the lines decreased and the differentiation between the lines increased. Therefore, the accuracy of genomic prediction between the lines (GBLUP-B) is very low (0.05) and there is a small increase in accuracy when adding data from other lines (GBLUP-C) to the training population (Figure 5). Moreover, across the 10 replicates around 30% of the GBLUP-B values in each cycle are negative. A pattern of reproduction without gene flow between the populations results in the production of divergent breeding populations (lines) with high genetic distances between them (as a result of the isolation). After many breeding cycles, this isolation could lead to differentiation between the populations resulting in major changes in the strain’s characteristics and develop one or more sub-strains in the breeding programme.

Allowing mixing between lines at a rate as low as 4%, we observe that the lines become genetically more similar as the genetic distance between them decreases, while they have more genetic variance as the within line genetic distance increases over the 10 cycles. This results in an increase (≈10%) in the accuracy of the prediction by combining all the populations and therefore a gain of using a larger training population for the evaluation, instead of using data only for one line to predict GEBVs of another line. At cycle 10, the GBLUP-C scheme has 5.3% higher values than GBLUP-W. By increasing further the mixing rate, the number of cycles that is needed to make the lines more similar reduces and the genetic diversity within them increases. With high mixing rates, the decrease of the genetic distance across the lines is rapid but the non-consecutive lines need more cycles in order to become as similar. Consequently, the prediction accuracy from one line to another (GBLUP-B) increased rapidly (>50%) over the 10 cycles. Moreover, the gain in accuracy is higher when using data from all lines (GBLUP-C) compared to GBLUP-W and is observed sooner with higher rates of mixing. At cycle 10, the GBLUP-C scheme has more than 10% higher accuracy than GBLUP-W from mixing schemes 20 to 50%.

Key factors for the accuracy of genomic prediction are the relationships between the reference and target populations (the lines in our case) and the size of the reference population (jointly combined lines). In theory, by increasing the size of the reference population we expect an increase in the accuracy of genomic prediction (Goddard and Hayes 2009). Combining populations, in order to increase the reference population and therefore improve the accuracy of genomic prediction, is very useful and common in terrestrial animals e.g. in dairy and beef cattle (VanRaden et al. 2009, Lund et al. 2011, Chen, Vinsky and Li 2015) or in pigs (Hulsegge et al. 2019, Song et al. 2019). Taking into consideration that aggregating a large reference population can be easier in aquaculture breeding programmes than terrestrials, as the former can have bigger population sizes, combining populations could be a very useful tool to increase the available information in order to maximise the accuracy of the estimated breeding values. Moreover, aquatic species can obtain higher response to selection and greater genetic improvements because of their higher fecundity which allow them to be subject to much higher selection intensities in their breeding programmes.

A mixing rate of 30% or above seems an efficient mixing strategy for an aquaculture breeding programme. After only two breeding cycles, there is higher genetic distance within a population (which is equivalent to an increase in the genetic diversity available in the population) and at the same time higher genetic similarity between the populations (decreased between-line genetic distance). This results in an increased accuracy of genomic predictions for all tested schemes. After three breeding cycles the GBLUP-C accuracy is more than 10% higher than GBLUP-W for mixing rates 30 –50% and the GBLUP-B accuracy keep increasing for each cycle, even when the lines are very similar after only two cycles.

A typical Atlantic salmon breeding programme is based on different family–based schemes in which the trait of interest is tested on the sibs of the candidates. Subsequently, test information is used to calculate the breeding values for the selection of the parents (Gjedrem 1985). Unfortunately, only half of the genetic variation, that coming from the between family variation, has a use in these schemes. Thus, these schemes are used for traits that cannot be measured on the selection candidates such as disease challenge testing and carcass quality traits. The operation and maintenance of different sib-test population is very difficult and costly for the aquaculture companies. As parents cannot be reused across the generations, it is very important to find new strategies to reuse the parents’ information or to mix parents from different generations. This can be achieved either by using cryopreservation methods (Figueroa et al. 2016) or using artificially early smoltification individuals (Handeland et al. 2013) to increase the connectedness of the breeding lines. Unfortunately, the problem with these early smolts is that a low proportion of them become sexually mature at three years of age. This makes it more difficult to obtain the necessary number of breeders that the breeding programme requires. A further investment on artificial light technologies which simulate different photoperiods will increase this percentage of mature fish at three years of age.

For practical aquaculture breeding programmes, the results of this simulation study could have many implications, which could lead to modifications in existing breeding programmes. Breeding companies could reduce phenotyping costs by increasing connectedness between populations through mixing as this would allow to reduce (or eliminate) sib-testing of the candidates since GBLUP-C could provide more accurate GEBVs on populations with no available data. As many salmon breeding programmes are family-based, the most important traits are actually measured on the sibs of the candidates and then the test information is used to select the best candidates by estimating their breeding values (Gjedrem 1985). This phenotypic testing must be repeated for each breeding line and for each trait of interest (disease challenges or slaughter phenotypes), which constitute a large part of the cost of the breeding programme. By increasing the connectedness of the breeding lines (mixing) and using jointly all the available data (GBLUP-C scheme) there is no need to repeat this process for each line. Thus, the GEBVs can be estimated with higher accuracy between the lines (GBLUP-W scheme), which results in improvement in the accuracy of candidate selection, reducing the number of sib-tests and improving animal welfare.

In conclusion, our results show that by increasing the mixing rate, the populations become more similar, so the information of other populations are useful in the evaluation and thus the GBLUP-B and GBLUP-C lead to greater improvement in the accuracy. This could yield immediate benefits in aquaculture breeding programmes as increasing the similarity of breeding lines will improve the selection of candidates, reduce the breeding cost (by reducing sib-tests and reuse of genotype data) and also improve animal welfare. In this study, we demonstrate how accuracy can be increased with different mixing rate under a random selection breeding programme. Thus, future studies will be required to investigate how different mixing rates can affect the genetic gain and the accuracy of GEBVs in a directional selection breeding programme.

## Availability of data and materials

The dataset used in the current study is available from the corresponding author on reasonable request.

