## Supplementary Material for "Impact of mixing between parallel year groups on genomic prediction in Atlantic salmon breeding programmes under random selection"

**Table A1. The overall average accuracy of genomic prediction with different mixing rates through the 10 cycles. The “Rate” is the proportion of mixing individuals from one line to another. The “GBLUP-W” model use only data of the same population, the “GBLUP-B” using data of one line to calculate the GEBVs for the individuals of another population and “GBLUP-C” using all data jointly to calculate GEBVs of individuals of one population.**

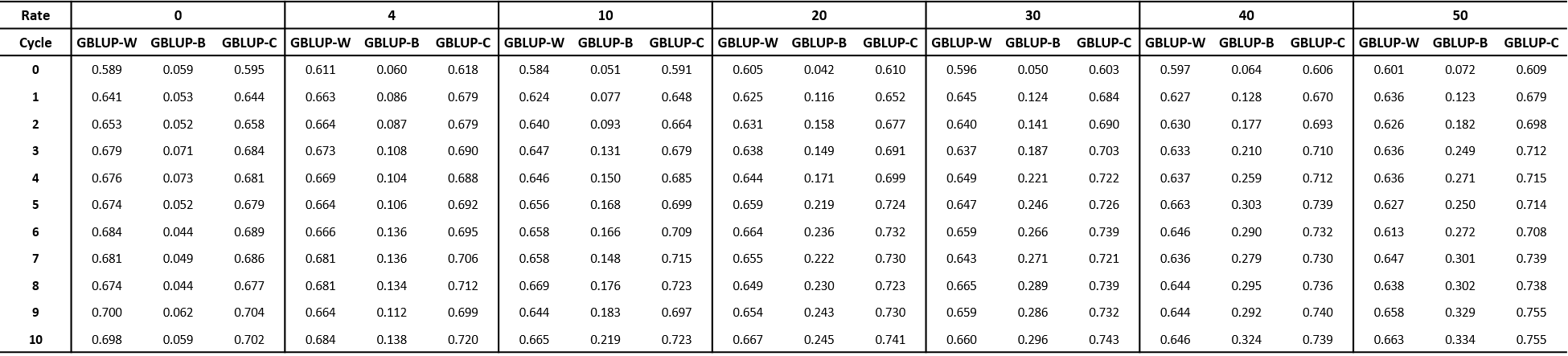

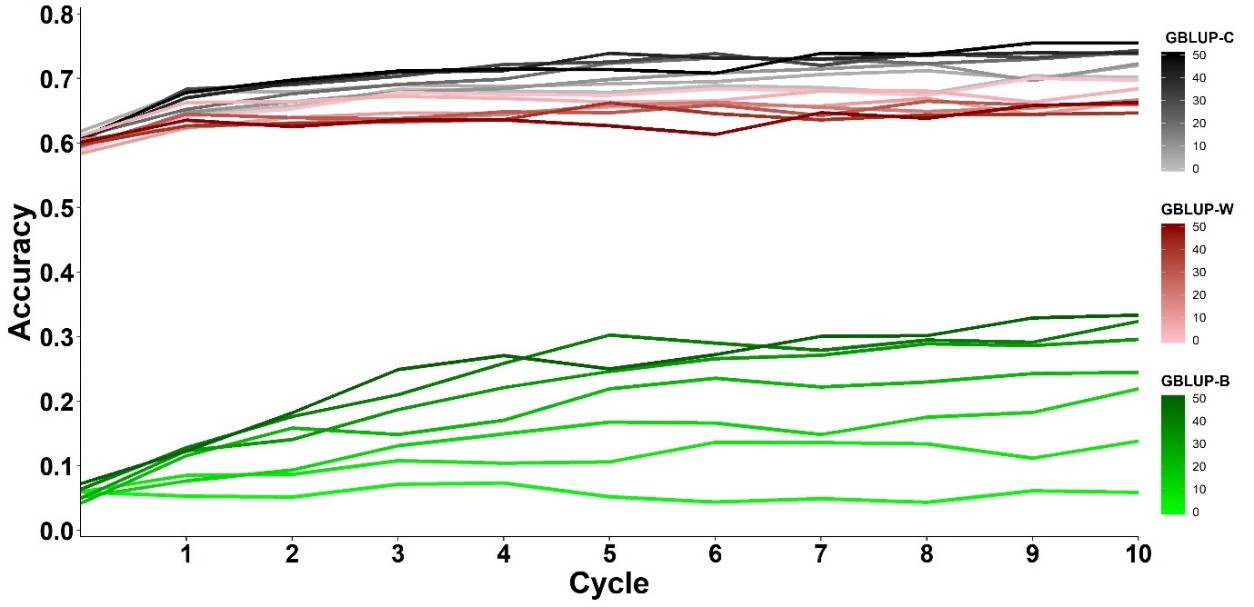

Figure A1. Different types of accuracy of genomic prediction across all the mixing rate scenario using data with the three following ways: “GBLUP-W”, “GBLUP-B” and “GBLUP-C”. Results are presented for three different schemes: A) Using data from only one line to estimate the GEBVs of the same line (GBLUP-W), B) using data of two lines where in one line the data are known and we estimate the accuracy of predicting GEBV of the other line (GBLUP-B) and C) using data from all the populations jointly to estimate the accuracy of predicting GEBVs of one line (GBLUP-C).
